# Gene dosage effects of polyA track engineered hypomorphs

**DOI:** 10.1101/2021.02.10.430645

**Authors:** Geralle Powell, Slavica Pavlovic-Djuranovic, Sergej Djuranovic

## Abstract

The manipulation of gene activity through the creation of hypomorphic mutants has been a long-standing tool in examining gene function. Our previous studies have indicated that hypomorphic mutants could be created by inserting cis-regulatory sequences composed of consecutive adenosine nucleotides called polyA tracks. Here we use polyA tracks to create hypomorphic mutants and functional characterization of membrane, secretory and endogenous proteins. Insertion of polyA tracks into the sequences of interleukin-2 and membrane protein CD20 results in a programmable reduction of mRNA stability and attenuation of protein expression regardless of the presence of signaling sequence. Likewise, CRISPR/Cas9 targeted insertion of polyA tracks in the coding sequence of endogenous human genes *AUF1* and *TP53* results in a programmable reduction of targeted protein and mRNA levels. Functional analyses of AUF1 engineered hypomorphs indicate a direct correlation between *AUF1* gene levels and the stability of AUF1-regulated mRNAs. Hypomorphs of TP53 affect the expression of the target genes differentially depending upon the severity of the hypomorphic mutation. Finally, decreases in TP53 protein affect the same cellular pathways in polyA track engineered cells as in cancer cells, indicating these variants’ biological relevance. These results highlight this technology’s power to create predictable, stable hypomorphs in recombinant or endogenous genes in combination with CRISPR/Cas9 engineering tools.

## INTRODUCTION

The investigation of a gene’s phenotype and its functional importance has often occurred by creating loss-of-function (LOF) mutations. LOF mutations allow the examination of phenotype when gene expression is reduced (as with hypomorphic mutations) or completely ablated (as with “null” mutations).^1^ However, phenotypic investigation of LOF mutations can be troublesome due to embryonic lethality or disruptions in organismal development as a result of pleiotropy.^2,3^ Therefore, the use of hypomorphic mutations is often preferable and more experimentally advantageous.

There are several methods for creating hypomorphic mutations. Historically, chemical and UV irradiation mutagenesis screens followed by selection for phenotypic strength have been used to identify hypomorphic alleles.^1,4–6^ Hypomorphic mutations have also been created through non-directed insertional mutagenesis using transposable elements in or near regulatory regions.^7,8^ RNAi has also been an essential tool in creating hypomorphic phenotypes due to varying degrees of residual gene expression produced from knockdowns.^9–11^ However, each method for the creation of hypomorphic mutations is accompanied by its disadvantages. Although the advance in the next-generation sequencing has reduced much of the time and tediousness associated with finding phenotypic variants in traditional forward genetics screens, the process still requires creating a mapping population, with mapping resolution being variable.^12–14^ Even if an allele is mapped to a gene of interest, mutagenesis screens also produce many other types of alleles, such as neomorphs (altered or unique function), nulls (loss-of-function), and hypermorphs (gain-of-function). On the other hand, RNAi results in the variable gene knockdown and can have off-target effects.^15^ As such, there is a need for a method that can be used for all recombinant coding genes and, in combination with novel CRISPR/Cas9 technologies, to engineer endogenous genes.

We have previously described polyA tracks as stretches of adenosine nucleotides that decrease mRNA stability during the elongation phase of mRNA translation, causing ribosomal stalling and frameshifting.^16^ The direct consequence of these events is a decrease in protein expression and mRNA stability.^16–20^ The decrease in the amount of gene expression directly correlates to the length of the polyA track: the longer the polyA track, the lower the mRNA’s stability and the expression of the protein products.^16,17,19,20^ Our previous work indicated that polyA tracks could be used to create hypomorphic levels of recombinant cytoplasmic proteins in various reporters and across multiple organisms such as *E*.*coli, S. cerevisiae, T. thermophile, N. benthamiana*, and within Drosophila and human cells.^19^ Additional studies have recently indicated that the same technology can be used in *C. albicans* and *A. thaliana* cells, where constructs with polyA tracks were inserted into previously engineered knock-out cells. ^20^

The previous studies of polyA track hypomorphs have been limited to reporters or recombinant cytoplasmic proteins and were not tested in secretory or membrane proteins.^19^ Because the sequence and structure of the N-terminus of secretory and transmembrane proteins are essential for co-translational translocation and thus proper processing and transport, it remains unclear whether the insertion of polyA tracks would create hypomorphic mutants of such genes. ^21–23^ Moreover, it is still unknown whether polyA tracks could be engineered in endogenous genes by using the CRISPR/Cas9 system.

In this work, we show how the insertion of the polyA tracks into genes for integral membrane-spanning 4A1 (further referred to in this study as CD20) and secretory IL2 (Interleukin-2) regulates their recombinant protein expression. Inserting a polyA track of at least 9 and up to 18 adenosines reduces mRNA stability and protein synthesis in a programmable without impacting protein membrane insertion or secretion out of the cells. This effect is also independent of the presence of the signaling peptide sequence in IL2, where insertion of a polyA track immediately before and after the signal sequence results in similar protein and mRNA reduction. We show that the same effects can be observed when polyA track sequences are introduced into endogenous genes in human tissue cultures using CRISPR/Cas9 technology. AUF1 (AU-rich element RNA-Binding Protein 1) and TP53 (tumor protein p53) mRNA stability and protein expression are reduced depending on the length of polyA tracks in a programmable way. Analyses of downstream effects of AUF1 hypomorphs indicate changes in abundance of MAT1A (methionine adenosyltransferase 1A), APP (amyloid beta precursor protein), TOP2A (DNA topoisomerase II alpha), and USP1 (ubiquitin specific peptidase 1) mRNAs, previously shown to respond to AUF1 protein. ^24,25^ Inserting polyA tracks in the TP53 locus of HAP1 cells leads to the partial or near-complete loss of TP53 protein and TP53 dosage-dependent effects on multiple TP53-regulated genes. Our data also indicates that either partial or more complete loss of *TP53* gene expression leads to downregulation of *MGMT* (O-6-methylguanine-DNA methyltransferase) gene products, a trend observed in engineered cells and cancer cell lines with compromised levels of TP53 protein.

Overall, the polyA track method successfully creates hypomorphic mutations in secretory and membrane proteins as well as in endogenous genes. We also show the successful application of this method in both haploid and diploid genomes. Finally, the creation of hypomorphic mutants using CRISPR/Cas9 in conjunction with polyA track sequence insertion allows for the controlled investigation of downstream genes and pathways. This method provides titratable gene expression in a feasible and experimentally tractable way.

## RESULTS

### Insertion of polyA tracks creates hypomorphic mutants of membrane gene CD20

To investigate whether polyA track technology can be used for membrane proteins, we focused on CD20. ^26^ CD20 is a B lymphocyte-specific integral membrane protein essential for the differentiation of these cells. ^27^ As a member of the membrane-spanning 4A gene family encoded by the MS4A1 gene, CD20 is a 33-37 kDa protein with four hydrophobic transmembrane helices, one intracellular loop, and two extracellular loops with cytosolic N and C termini, respectively.^28^ Because CD20 lacks an N-terminal signal peptide, its correct expression and translocation depend on its first transmembrane domain.^27–29^

To test whether insertion of polyA tracks can lead to programmable expression of CD20, we cloned the full sequence of CD20 into the pDEST40 vector and inserted polyA tracks of 12 adenosines (12As, sequence equivalent to 4 consecutive lysine AAA codons) and 18As (sequence equivalent to 6 lysine AAA codons), respectively, after the second amino acid (Fig. 1 A). We transfected WT and mutated CD20 reporters together with a GFP-YFP reporter as a normalization control into Chinese hamster ovary (CHO) cells. Using qRT-PCR and Western blot analysis, we examined the effects of polyA insertion on CD20 mRNA stability and protein expression (Fig. 1B and 1C). The recombinant CD20 protein was visualized using polyclonal CD20 antibody and compared to the expression of GFP-YFP constructs. The expression of CD20 protein was decreased over approximately 50% and 80% for reporters with the insertion of 12 and 18As, respectively (Fig. 1B) with previously observed degradation bands at 25kDa.^29^ We observed a similar approximate 50% and 70% reduction in mRNA stability for constructs with the insertion of 12 and 18As, respectively (Fig. 1C).

**Figure 1.**
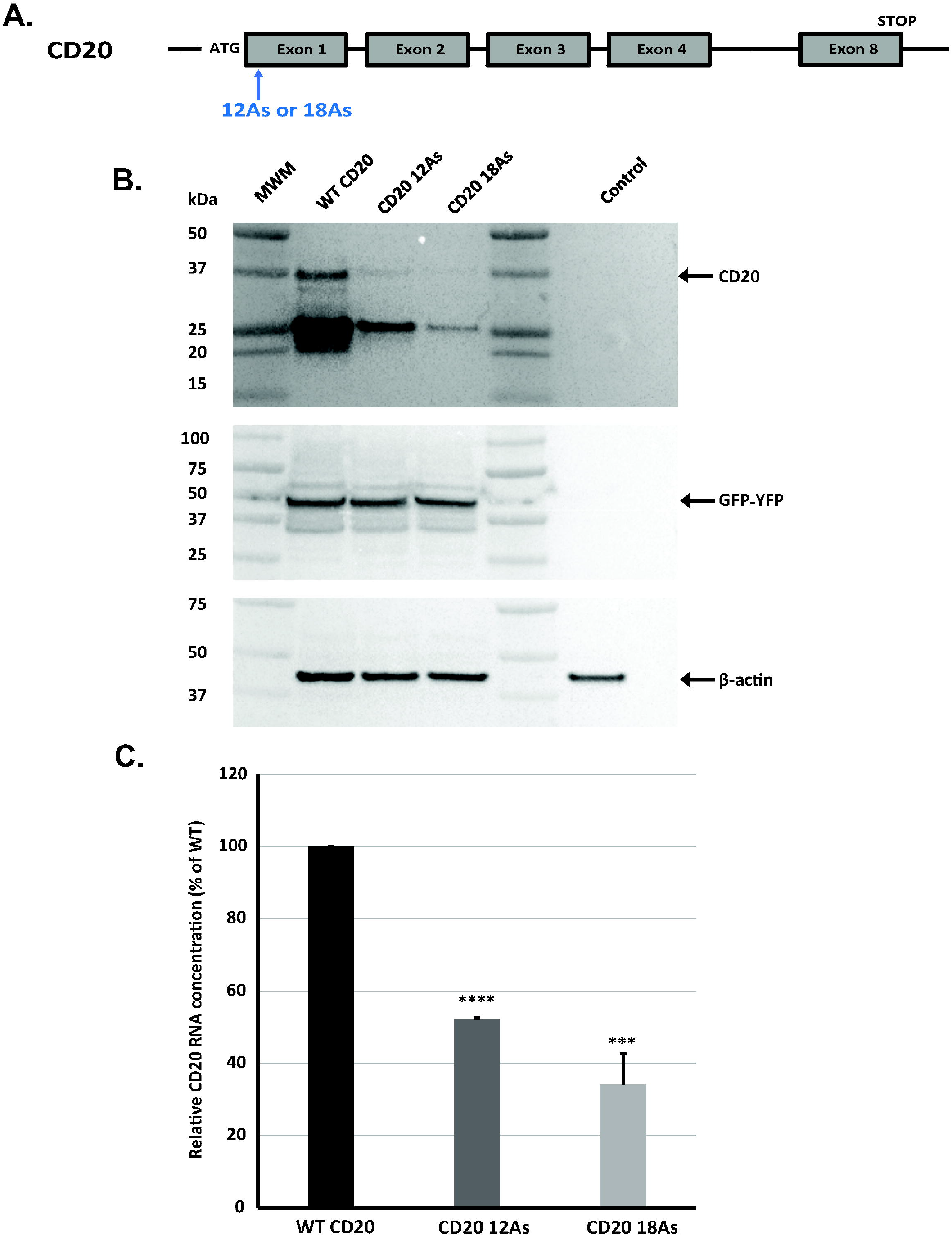
Design and effect of polyA track insertion on CD20 stability and expression. A. Scheme representative of polyA insertions and location of insertions for CD20 reporter constructs used in this study. PolyA insertions in the CD20 coding sequence were placed proximal to the ATG start codon. Schematic of *CD20* gene with start codon (ATG), termination codon (STOP) and exons and insertion site are indicated. **B**. Western blot analysis of CD20 constructs expressed transiently in CHO cells. Equal amount of total protein was used for analysis. GFP-YFP and β-actin were used as transfection and loading controls, respectively, with expression of both proteins detected using specific antibodies. CD20 was detected with CD20-specfic antibody. Arrows indicate position of full length CD20, GFP-YFP and β-actin proteins. **C**. mRNA levels of CD20 constructs were measured by qRT-PCR. CD20 expression was normalized to B-actin. Relative mRNA levels for both (12As) and (18As) constructs are presented as percentage of wild-type CD20. Error bars represent mean±s.d. values (*n*=3). P-values represent signficance (P ≤ 0.05,*;P ≤ 0.01, **; P ≤ 0.001,***; P ≤ 0.0001, ****).

To investigate whether insertion of polyA tracks and expression of CD20 protein with short poly-lysine repeats interferes with cellular localization of this membrane integral protein, we used live-cell imaging (Fig. 2A) and flow cytometry analysis (Fig. 2B). Using FITC-labeled CD20-2H7 specific antibody, we performed live-cell imaging of CHO cells expressing wild-type CD20 and polyA track variants. The monoclonal CD20 antibody 2H7 binds an epitope found in the larger extracellular loop of CD20 and has been used for the depletion of B-cells.^30,31^ Antibody immunofluorescence indicated that all three variants of CD20 are expressed in the membrane of CHO cells.^32^ Furthermore, the position of the FITC-labeled CD20-2H7 specific antibody indicated the distribution of CD20 similar to the CellMask™ Orange plasma membrane stain (Fig. 2A and Supplementary Fig. 1). The difference in the levels of cellular immunofluorescence was similar to the difference in protein levels seen in western blot analyses (Fig. 1B). Cells expressing wild-type CD20 construct had the highest immunofluorescence signal, while polyA track engineered constructs had a programmable reduction in the abundance of the CD20 protein. CD20 protein with insertion of polyA track of 12As had higher immunofluorescence than CD20 with 18As. Flow cytometry of transfected CHO cells labeled with FITC-labeled CD20-antibody also indicated detectable levels of expression of wild-type CD20 and CD20 with polyA track of 12As (Fig. 2B) .^33^

**Figure 2.**
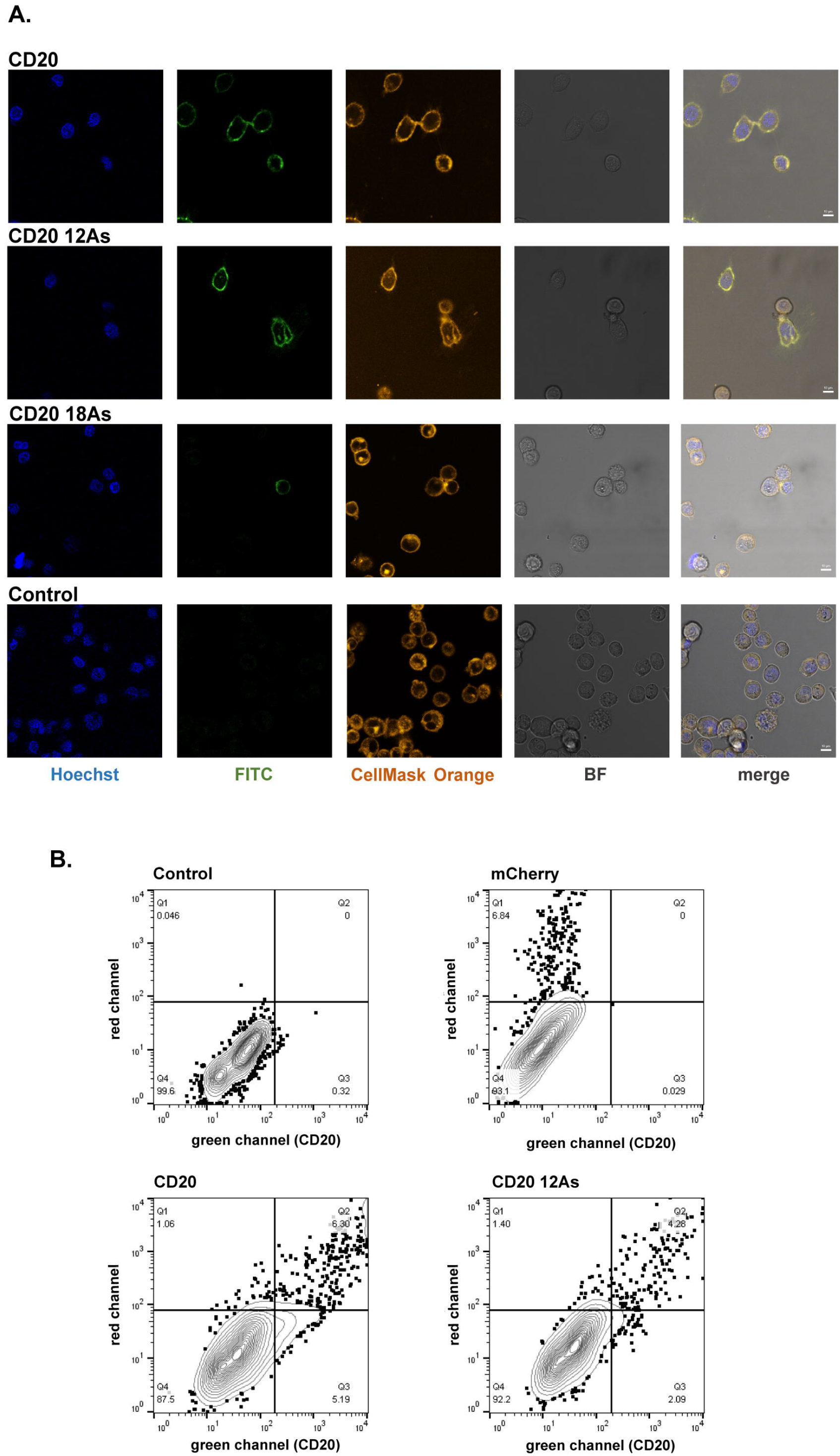
CD20 hypomorphs live imaging and flow cytometry. **A**. Fluorecent and bright field (BF) microscopy images of live CHO cells expressing CD20, CD20 12As and CD18As constructs. CD20 positive cells were labelled using anti-CD20-2H7 FITC-labelled antibody. The cell membrane is labeled with CellMask™ Orange stain. The control non-transfected CHO cells are indicated. The DNA was labeled with Hoechst 33342 stain. Scale bar represents 10 um. **B**. Representative graphs showing flow cytometry scatter plots of CHO cells expressing CD20 and CD20 12As co-transfected with mCherry constract. CHO cells expressing only mCherry and non-transfected cells are used as controls. The cells were stained with anti-CD20-2H7 FITC-labelled antibody.

### Insertion of polyA tracks creates hypomorphic mutants of secretory IL2 protein regardless of signaling peptide

To test the ability to create hypomorphic mutants using polyA tracks in secretory proteins, we created reporter genes with the full-length sequence of IL2. IL2 is a cytokine necessary for the homeostasis of T-lymphocytes and the clonal expansion of regulatory T-cells.^34,35^ As a secretory protein, IL2 contains an N-terminal signal peptide for recognition by the SRP and efficient co-translational translocation. It was shown previously that modification of the signal peptide of IL2 could affect IL2 expression and secretion.^36^

To test the locational flexibility of polyA insertion, we placed polyA tracks of 9, 12, and 18As, respectively, before and after the signal peptide sequence (Fig. 3A). PolyA tracks before the signal peptide were inserted after the codon for the second amino acid, while insertion after the signal peptide sequence occurred following the codon for amino acid 27. We transfected WT IL2 and IL2 reporters with polyA track insertions into Human embryonic kidney 293 (Hek293) cells. These cells were previously shown to produce secreted IL2 protein when transfected with recombinant IL2.^37,38^ We assayed mRNA amounts by qRT-PCR analysis for all reporters. RNA levels were reduced by approximately 20, 40, and 50% for 9, 12, and 18As, respectively, for polyA tracks inserted before the signal sequence and 30, 50, and 70% for polyA tracks inserted after the signal sequence, respectively (Fig. 3B). We used cell media to measure the amount of secreted IL2 protein after 48 hours after transfection and applied it to a standardized IL2-sandwich ELISA (Thermo Fisher, EH2IL2). To check the linearity of the IL2-ELISA test, we diluted wild-type IL2 samples from 2 and up to 16 fold and used media from untransfected cells as control (WT NI). Similar to the mRNA experiments, protein expression was reduced in IL2 variants with polyA tracks regardless of position respective to the signal peptide sequence (Fig. 3C). Insertion of 9As, 12As or 18As, consistently led to 60-70, 80, or more than 90% reduction in the IL2 levels as detected by an ELISA assay (Fig. 3C), regardless of whether insertion of the polyA track was before or after signal peptide.

**Figure 3.**
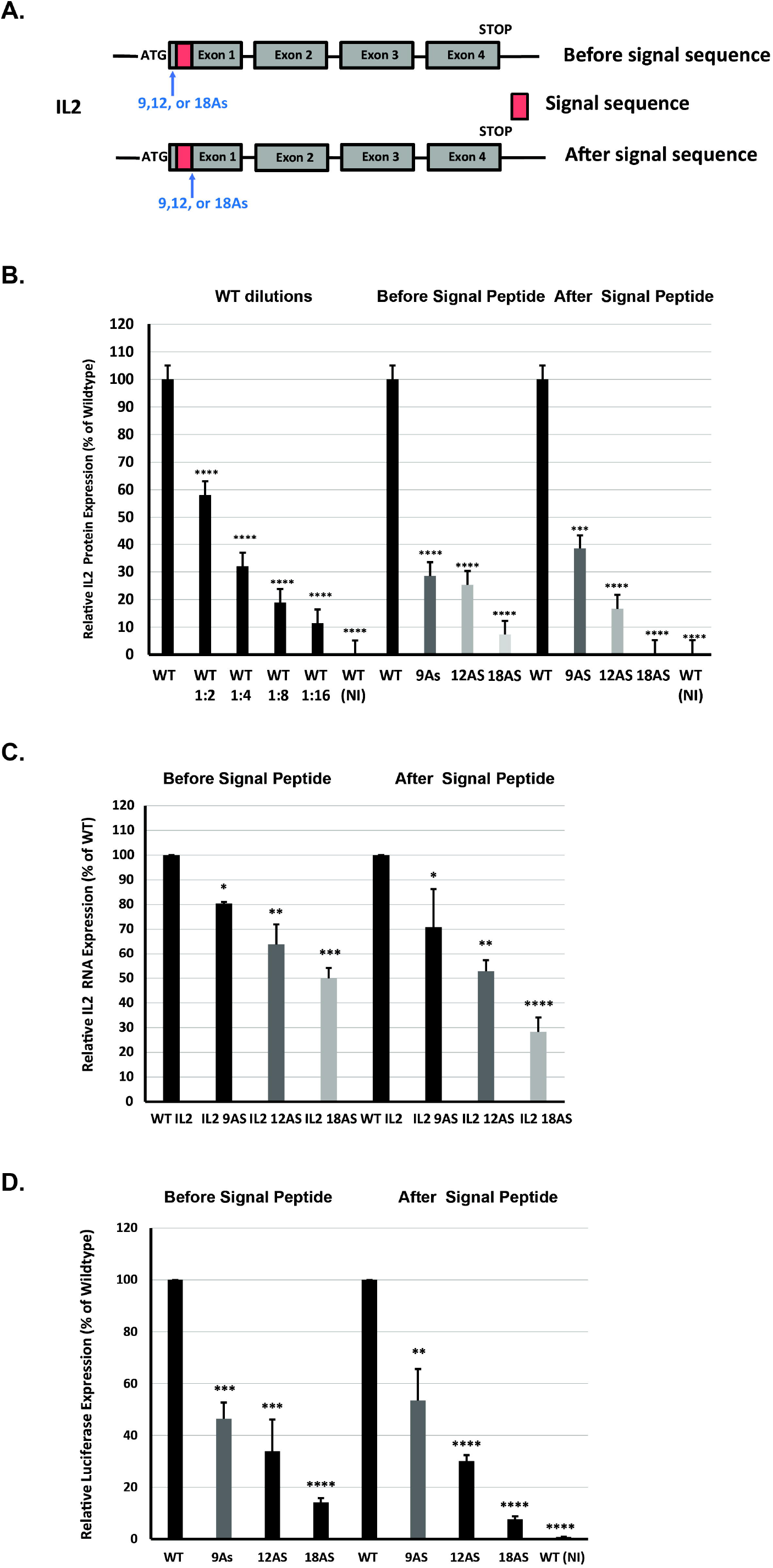
Effects of polyA insertions on IL2 reporter expression, stability and activity. **A**. Scheme representative of polyA insertions and location of insertions for IL2 reporter genes used in this study. PolyA insertions were placed before and after the signaling sequence (SS). IL2 constructs containing full-length wild-type IL2 sequence (WT IL2), and insertions of 9A, 12A, and 18As, respectively, were transiently expressed in HEK293 cells. Schematic of IL2 gene with start codon (ATG), termination codon (STOP), signal sequence, exons and insertion sites are indicated. **B**. ELISA measured levels of IL2 protein expression. IL2 protein levels are calculated from cell supernatant collected from control (WT NI) HEK 293 cells and cells expressing WT IL2 and IL2with polyA tracks (9, 12 and 18As), respectively. Standard curve was used to get mean net absorbance. Dilutions of the supernatants from WT IL2 (1:2 – 1:16) indicate linearity of the assay. **C**. mRNA levels of IL2 constructs were measured by qRT-PCR. IL2 expression was normalized to β-actin. Relative mRNA and protein levels for control (-), WT IL2, and IL2 polyA tracks constructs (9, 12 and 18As) are presented as percentage of wild-type IL2. **D**. Biological activity of IL2 measured by IL2 Bioassay (Promega). Relative IL2 activity is measured by luciferase expression and normalized to supernatant of WT IL2 construct. Control (WT NI) and IL2 polyA tracks constructs (9, 12 and 18As) are presented as percentage of wild-type IL2. Error bars represent mean ± s.d. values (*n*=3). P-values represent signficance (P ≤ 0.05,*;P ≤ 0.01, **; P ≤ 0.001,***; P ≤ 0.0001, ****).

Finally, to test the functionality of the IL2 variants with polyA tracks, we used IL-2 cell-based assay (IL2 Bioassay Promega), which reports the stimulation of IL2-receptor through the expression of luciferase reporter (Fig. 3D). Results for supernatants of either wild-type IL2 sample or IL2 variants with polyA tracks indicated functionally active proteins, regardless of the position of polyA track to encoded signaling peptide. Moreover, the luminescence levels from the functional IL2 assay indicated a similar change in the levels of IL2 protein and polyA track hypomorphs that we previously observed in ELISA assays (Fig. 3B).

### CRISPR/Cas9 insertion of polyA tracks in TP53 endogenous locus in near-haploid HAP1 cell line

With CRISPR/Cas9 technology development, a range of gene modifications, including point mutations, insertions, and deletions, can be made at precise genomic locations.^39^ We asked if endogenous hypomorphic mutants could be created using CRISPR/Cas9 technology to insert polyA track of various lengths in endogenous gene loci in human tissue cultures. Using this method, we created endogenous hypomorphic mutants in the *TP53* (tumor protein 53) gene and the *AUF1* (AU-rich element RNA-binding protein 1) in HAP1 and Hek293 cells, respectively (Fig. 4 and 5).

**Figure 4.**
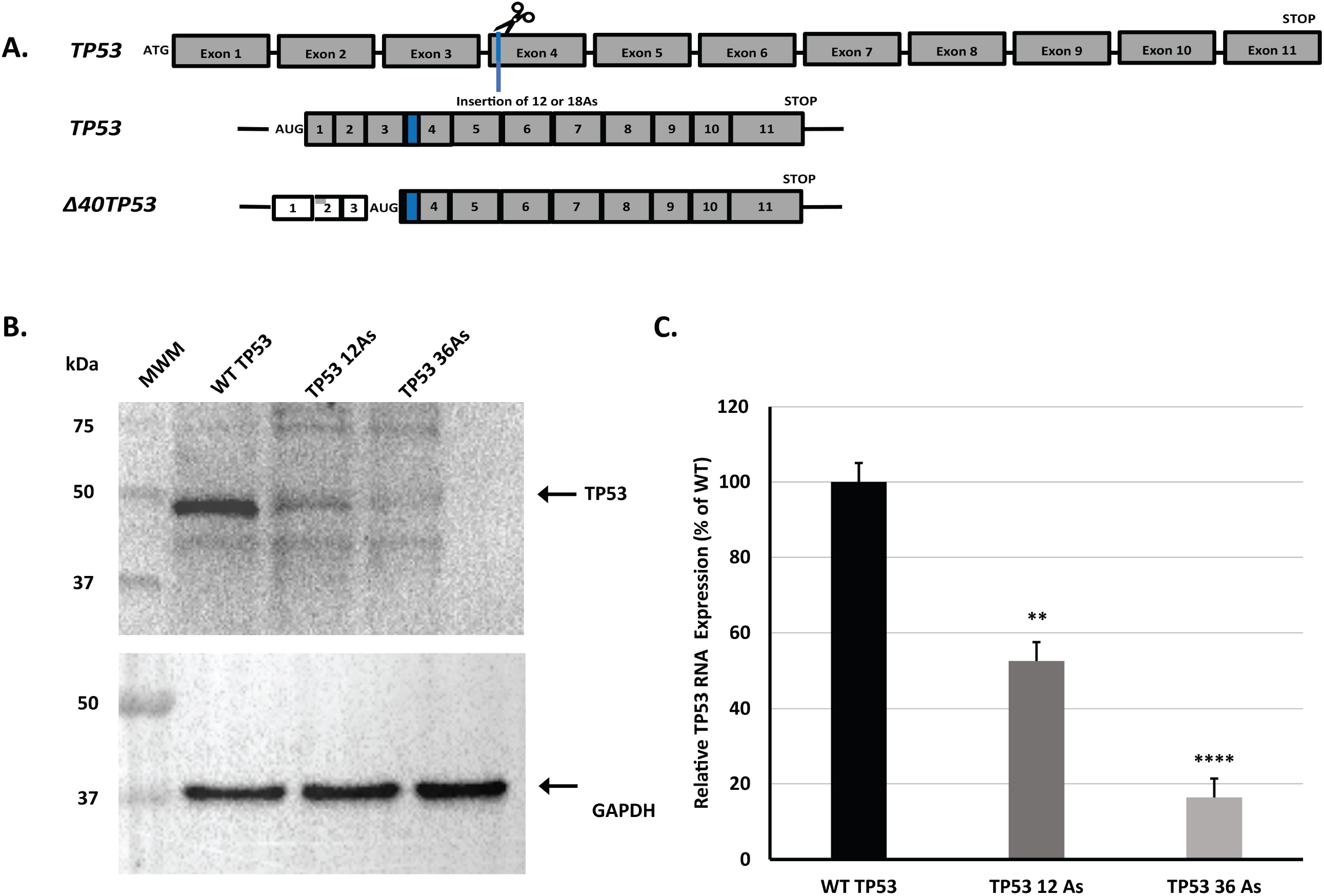
Creation and expression levels of endogenous TP53-polyA track hypomorphs in HAP1 cells using CRISPR/Cas9 genomic editing. **A**. Cartoon representing the design of CRISPR/Cas9 mediated insertion of polyA tracks in the DNA sequence of TP53. The blue dashed line indicates the Cas9 (scissors) cleavage site and location of polyA track insertion (12 or 36As). Both the canonical and N-terminally truncated (delta 40) isoforms are shown with the engineered polyA tracks (shown in blue). **B**. Western blot analysis of WT HAP1 cells and HAP1 cells with endogenous polyA track insertions of 12 and 36 consecutive adenosine nucleotides, respectively. Equal protein was used for analysis. TP53 was detected using TP53 specific antibody. Constitutively expressed GAPDH was used as a loading control and was detected using specific antibody. **C**. The mRNA level of TP53 in WT HAP1 and TP53 hypomorphs was detected by qRT-PCR. TP53 expression was normalized to GAPDH mRNA expression. Relative mRNAs levels are presented as a percentage of WT TP53. Error bars represent mean ± s.d. values (*n*=3). P-values represent signficance (P ≤ 0.05,*;P ≤ 0.01, **; P ≤ 0.001,***; P ≤ 0.0001, ****). Arrows indicate position of the full length proteins.

**Figure 5.**
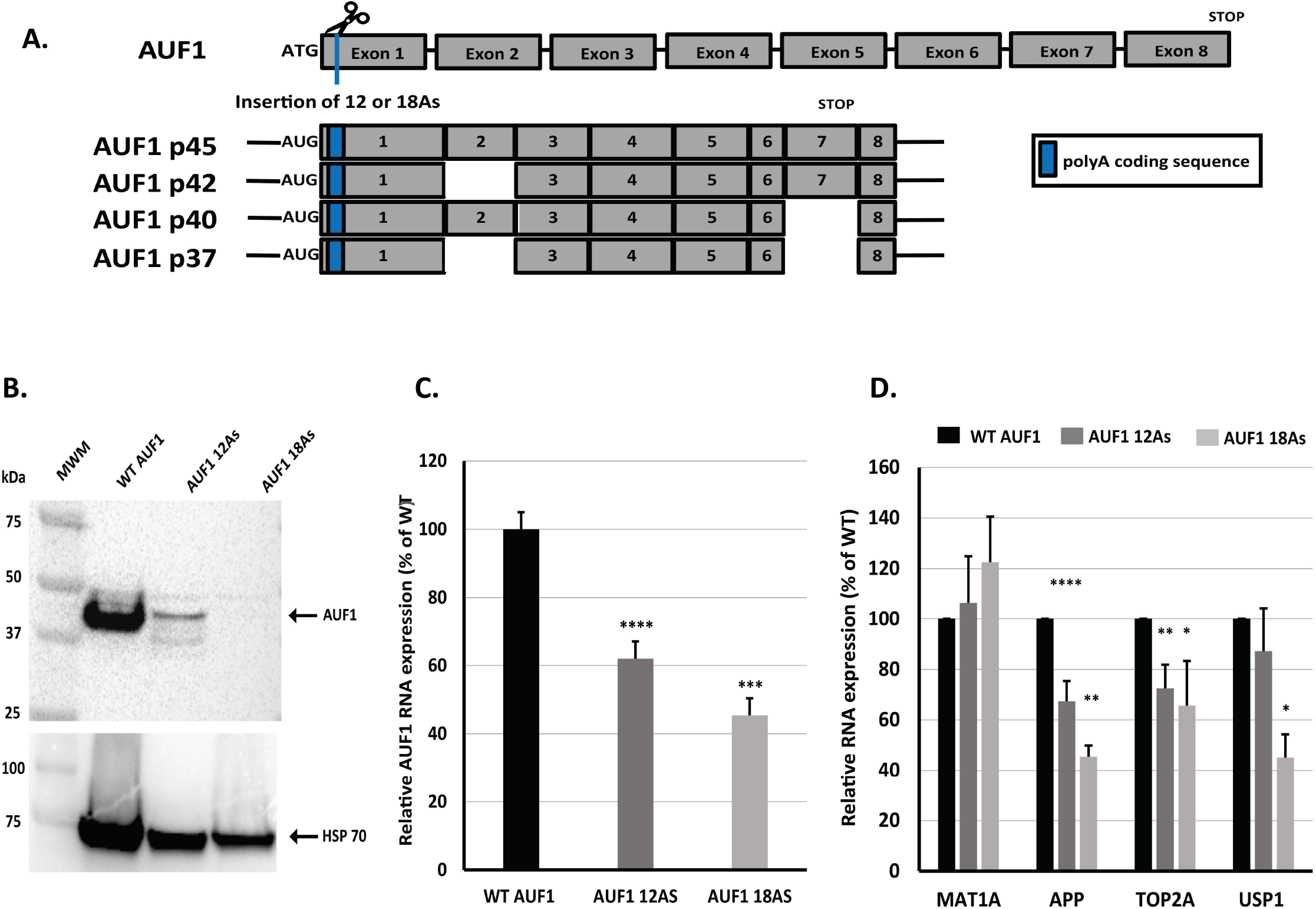
Design and gene-regulatory effect of endogenous AUF1 polyA track hypomorphs created using CRISPR/Cas9-genome engineering in HEK293 cells. **A**. Cartoon representing the location of polyA track insertion in the DNA coding region of AUF1. The dashed blue line represents the Cas9 (scissors) cut site and location of polyA insertion (12 or 18As). Four of the major AUF1 mRNA isoforms (p45,p42,p40,p37) are shown with their respective polyA insertions. **B**. Western blot analysis of WT HEK293 cells and HEK293 cells with endogenous polyA insertions of 12 and 18 consecutive adenosine nucleotides, respectively. Equal protein was used for analysis. AUF1 was detected specific antibody. Constitutively expressed HSP70 was used as a loading control and was detected using specific antibody. **C**. The mRNA level of AUF1 in WT HEK293 cells and AUF1 hypomorphs was detected by qRT-PCR. GAPDH was used for normalization of total mRNA levels. Relative mRNA levels are presented as a percentage of WT AUF1 as expressed in WT HEK293 cells. **D**. The mRNA levels of AUF1 downstream targets, MAT1A, APP, TOP2A, and USP1, were detected by qRT-PCR. GAPDH was used to normalize total mRNA levels. Error bars represent mean ± s.d. values (*n*=3). P-values represent signficance (P ≤ 0.05,*;P ≤ 0.01, **; P ≤ 0.001,***; P ≤ 0.0001, ****). Arrows indicate position of the full length proteins.

gRNAs were designed to facilitate Cas9 nuclease activity in the 4^th^ exon of the *TP53* gene (Supplementary Fig 2). This gene locus encodes the loop region in the TP53 protein. We selected this position to assure targeting of the two major N-terminal isoforms of TP53 transcripts: canonical TP53, which contains the full N-terminal sequence, and Δ40TP53, which excludes the first 40 amino acids of the canonical TP53 isoform due to the presence of an internal start (AUG) codon (Fig. 4A and Supplementary Fig. 2).^40^ gRNAs were chosen for high NHEJ (non-homologous end joining) efficiency, low off-target effects, and proximity to the cut site. ssODN (single-strand oligo donor) DNA containing homology arms flanking both the 3’ and 5’ regions of the polyA track insertion sequence (12 or 36As) was used as a template for CRISPR-mediated HDR (homology-directed repair) in the human near-haploid cell line HAP1.^41^

After CRISPR/Cas9 engineering of the *TP53* locus, we selected and confirmed by both NGS (next-generation sequencing) and Sanger sequencing multiple cell clones containing the insertion of 12 or 36As (Supplementary Fig. 2). The engineered cell lines with the successful insertion of polyA tracks in the single copy of the *TP53* gene in HAP1 cells were analyzed for mRNA stability and protein expression of the *TP53* gene using qPCR and Western blot analysis. GAPDH protein and mRNA levels were used as normalization controls between parental and engineered cell lines. Similar to the CD20 and IL2 reporters, endogenous TP53 hypomorphs containing 12 or 36 As had decreased mRNA levels and protein abundance dependent on the length of the inserted polyA track (Fig. 4B and 4C). The 12As insertion reduced both mRNA and protein amounts by 50% compared to wild-type TP53 expression in the original HAP1 parental line. Insertion of the 36As resulted in the almost complete loss of the TP53 protein and a 90% reduction in mRNA levels compared to TP53 levels in the original non-engineered HAP1 cells.

### polyA tracks engineered in AUF1 endogenous loci of diploid HEK293 cells result in programmable hypomorphs

To test whether CRISPR/Cas9 approach can be used to create polyA track hypomorphs in a human-derived diploid cell line, we targeted the AUF1 gene loci in the HEK293 T-Rex cell line (Invitrogen). As with the *TP53* gene, we introduced polyA tracks of 12 or 18As in the open reading frame of AUF1 (Fig. 5 and Supplementary Fig. 3). This time we engineered polyA tracks in the first exon of the *AUF1* gene after the second amino acid to assure targeting of all potential isoforms of AUF1 protein (Fig. 5A).^42^ After isolation of cell clones and confirmation of the polyA track insertion (Supplementary Fig. 3), cell lysates were analyzed for AUF1 mRNA levels and protein expression (Fig. 5B and 5C). Similar to TP53, the mRNA stability and protein expression of endogenously engineered AUF1 hypomorphs were decreased in a length-dependent manner. Insertion of the polyA track of 12As led to a 50% and 40% reduction in protein and mRNA levels of AUF1, respectively. Insertion of 18As further reduced protein levels to approximately 10% of the parental cell line levels, while mRNA levels were reduced by 60%.

### Hypomorphs of AUF1 affect mRNA levels of downstream target mRNAs

To test whether created hypomorphs with polyA insertions endogenously could be used to study the function of the gene with regards to established downstream targets, we performed expression analysis on both TP53 and AUF1 hypomorphs (Fig. 5D and 6). We measured the steady-state levels of the MAT1A, APP, TOP2A, and USP1 mRNAs in the parental HEK293 T-Rex cell line and the lines with engineered AUF1 hypomorphs (Fig. 5D). MAT1A mRNA was previously stabilized by knockdown of AUF1,while the transient knockdown of AUF1 by siRNA transfection induces the destabilization of APP, TOP2A, and USP1 mRNAs (34). Our results confirm the observation of previous studies.^24,25^ MAT1A mRNA levels increase in the programmed hypomorphs of AUF1, while levels of APP, TOP2A, and USP1 decline with a reduction in AUF1 expression. Even more compelling, the change in target gene mRNA levels was dependent on the length of the polyA track insertion and the extent of AUF1 reduction. The insertion of 18As has a more pronounced effect on mRNA stability than 12As (Fig. 5D).

### Hypomorphs of TP53 affect downstream genes in a dose-dependent manner

To extend our studies on the potential use of programmable hypomorphs for gene dosage effects, we analyzed how the insertion of polyA tracks in the *TP53* gene and reduction of *TP53* gene products affects downstream pathways known to be controlled or connected to TP53 protein. Based on the previous studies and availability of antibodies, we selected BRCA1, DDIT4, DUSP6, MGMT, and P21 as TP53 associated genes.^43–49^ We analyzed cell lysates by qPCR and Western blot analysis (Fig. 6A and 6B). We used parental HAP1 cells and engineered HAP1 cells with 12A and 36A insertion in the TP53 coding sequence (Fig. 3). Tp53 protein and mRNA levels in these cells reflected wild-type levels (WT TP53), a 50% reduction in TP53 (TP53 12As), and nearly complete loss of TP53 protein (TP53 36As) (Fig. 4B and 4C, as well as Fig 6A and 6B). Western blot analyses indicated that BRCA1 levels did not change regardless of the TP53 protein levels (Fig. 6A). Protein levels of DUSP6 and DDIT4 showed differential behavior in the presence of the varying TP53 levels (Fig. 6A). A 50% reduction of TP53 protein levels caused by insertion of polyA track with 12As led to a significant reduction of DDIT4 and almost a complete loss of DUSP6 protein in those cells (HAP1 TP53 12As). Both DDIT4 and DUSP6 were either not affected or slightly increased in cells with almost complete loss of TP53 (HAP1 TP53 36As) when compared to WT HAP1 cells. Finally, MGMT and P21 protein levels were severely depleted in cells expressing TP53 hypomorphs and compared to the WT HAP1 cells. Analyses of the steady-state mRNA levels for all genes indicated the same trend as protein levels (Fig. 6B). BRCA1 mRNA levels were unaffected between cell lines with different TP53 levels. DDIT4 and DUSP6 mRNAs showed a significant drop in the levels with a 50% TP53 protein reduction and wild-type or higher levels of mRNAs in cells with almost complete loss of TP53. Finally, MGMT and P21 mRNA levels were severely depleted in both TP53 engineered cell lines. We further tested whether TP53 and P21 levels can be changed if we apply hydroxyurea to HAP1 cells as a cytotoxic and genotoxic drug (Fig. 6C). ^50^ We observed moderate increase in wild-type TP53 levels after 3 and after 24 hour treatment. Levels of TP53 levels with12A polyA track changed only after 24 hours and we didn’t observe detectable levels of TP53 with 36A insertion even after 24 hours of hydroxyurea treatment. The P21 levels in wild-type sample were induced after 3 hours of hydroxyurea treatment and reduced after 24 hours. Cells with 12A TP53 hypomorphic variants showed delayed induction of P21 levels only after 24 hour hydroxyurea treatment while cells with 36A TP53 varaint failed to show any induction of P21 even 24 hours after hydroxyurea treatment (Fig. 6C).

**Figure 6.**
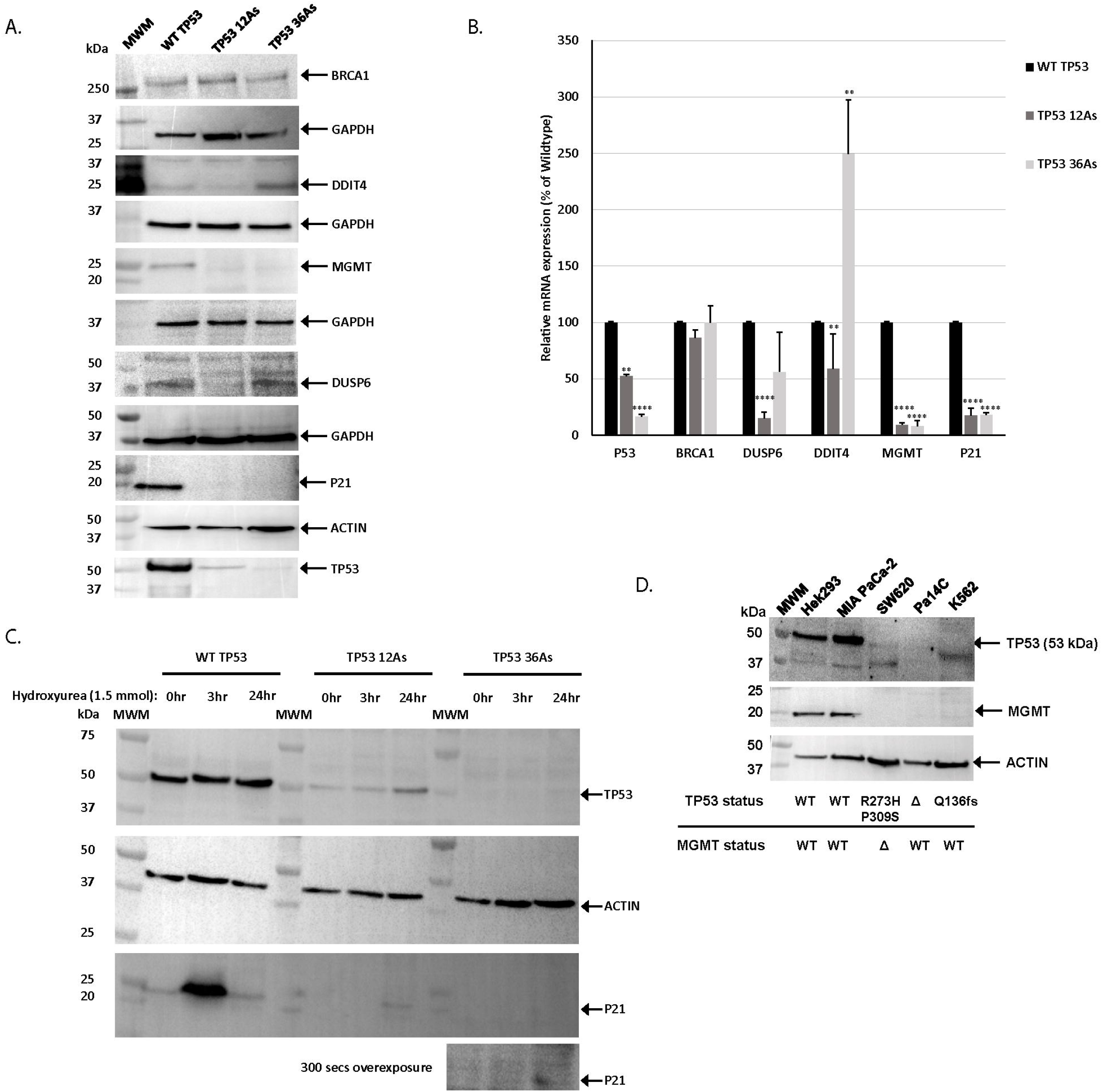
Effects of hypomorphic TP53 expression on the gene dosage of downstream targets. **A**. Western blot analysis of BRCA1, DDIT4, DUSP6, MGMT, P21 and TP53 protein levels in WT and TP53 polyA track (12 and 36As) HAP1 engineered cells. Equal amounts of total protein was used for analysis. BRCA1, DDIT4, DUSP6, MGMT, P21 and TP53 expression was identified using specific antibody. GAPDH and ACTIN were used a loading control in each blot and was detected using specific antibody. **B**. BRCA1, DDIT4, DUSP6, MGMT, P21 and TP53 mRNA levels in WT and TP53 polyA track (12 and 36As) engineered cells. Relative mRNAs levels are presented as a percentage of the mRNA levels of the wild-type gene as expressed in WT HAP1 cells. **C**. Western blot analyses of TP53 and P21 proteins in hydroxyrea treated and control (NT) cells. The length of hydroxyurea treatment is indicated for WT and TP53 polyA track (12 and 36As) engineered cells. **D**. Western blot analysis of TP53 and MGMT expression, respectively in HEK293, MIA PaCa2, SW620, Pa14C, and K562 cells. ACTIN was used to normalize total protein levels. Error bars represent mean ± s.d. values (*n*=3). TP53 and MGMT status are shown below (WT, wild type; Δ, null). P-values represent signficance (P ≤ 0.05,*;P ≤ 0.01, **; P ≤ 0.001,***; P ≤ 0.0001, ****). Arrows indicate position of the full length proteins.

Finally, to test whether the polyA track hypomorphs can reproduce some of the results of gene mutations or deletions found in the established cell from cancer patients, we obtained four TP53 mutant or deletion cell lines (Fig. 6D) and focused on MGMT protein levels. Based upon our hypomorphic models, we would predict that cell lines that contain less TP53 would contain less MGMT. In cell lines with wild-typeTP53 status (Hek293 and MIA PaCa-2), MGMT protein expression was easily detectible. The SW620 cell line has a double mutation in *TP53* gene (R273H/P309S) and a deletion of the MGMT gene. As expected, TP53 protein levels were greatly reduced in the SW620 cell line, and MGMT protein expression was not observed. In the two cell lines (PA14C (del) and K562 (Q136fs)) that have negligible amounts of TP53 protein, MGMT protein levels were completely depleted (Fig. 6D). These data indicate that our hypomorphic TP53 model can predict levels of the downstream gene MGMT.

## DISCUSSION

In this study, we have presented a comprehensive method of creating hypomorphic mutations through the insertion of polyA tracks. Using these powerful gene regulatory sequences, we show a method of creating hypomorphic mutations that is titratable and predictable (Fig. 1-4). We have expanded the use of polyA track insertions to create hypomorphic mutants in integral membrane proteins (Fig. 1). Integral membrane proteins make up 20-30% of the proteome with highly diverse functions. Integral membrane proteins constitute the lipid bilayer of the membrane surrounding cellular organelles and the outside of the cell. They thus modulate movements of molecules and information across membranes and serve essential functions as signal receptors and transporters of solutes. ^51,52^ Their importance to the cell function is apparent when these systems go awry; a substantial amount of disease-linked mutations occur in integral membrane proteins. Subsequently, these proteins make up over 50% of FDA approved drug targets.^53^

As a result of their significance in both health and disease, we chose to demonstrate the ability to create hypomorphic mutations in the transmembrane B cell antigen CD20 (Fig. 1). By inserting 12 or 18 adenosine nucleotides directly following the second amino acid, we observed that CD20 mRNA expression was reduced by 50 and 60%, respectively. Protein expression was similarly reduced for both polyA track-containing reporters. In addition to successfully reducing CD20 expression using this method, Western blot analysis using CD20-protein-specific antibody indicated expression of the full-length CD20 at the expected size of 33kd and previously observed degradation product at 25kDa.^29^ Furthermore our live cell imaging and flow cytometry (Fig. 2 and Supplementary Fig. 1) indicated correct membrane localization and programmable reduction of membrane inserted CD20 protein supporting our hypothesis that introducing additional adenosine nucleotides at the N-terminus of the CD20 would not further affect localization and quality of the protein.^54^ This result shows that the N-terminal addition of the polyA track in other membrane genes may successfulyl create hypomorphic mutants.

Like membrane proteins, secretory proteins make up a large percentage of the proteome (∼11%) and serve diverse functions as hormones, cytokines, and members of the extracellular matrix. ^36,55,56^ Because of these diverse functions, secretory proteins are essential to immune responses and cell-to-cell communication. Furthermore, the recent development of CAR-T cell therapies indicates a need for attenuation of specific cytokines.^57^ By creating hypomorphic mutations in alleles, these genes’ function can be examined or modulated, adding to the knowledge and significance of these genes in health and disease. With the insertion of polyA tracks in these secretory genes, we wanted to ensure that these additions or mutations would not affect protein secretion by affecting the signal peptide’s proteolytic cleavage. We used the cytokine IL2 as a candidate due to the presence of a signal peptide at its N-terminus (Fig. 3).

To test the polyA tracks’ placement on IL2 RNA stability and secretion, we created gene reporters containing the complete coding sequence of IL2 with polyA tracks of 9,12, or 18 adenosine nucleotides inserted before or after the signal peptide, respectively (Fig. 3A). Placement of the polyA track in IL2 did not affect IL2 secretion, as shown by the measurable amount and functionality of IL2 protein found in the supernatant of transfected HEK 293 cells (Fig. 3B and 3D). The mRNA stability of *IL2* and its protein expression was decreased for all constructs, irrespective of the location of the polyA track insertion. Furthermore, as shown in our previous studies in which we created hypomorphic reporter genes in eukaryotic cells, the decrease in IL2 gene expression is titratable, meaning the gene knockdown depends on its length the polyA track.^19^ While the magnitude of IL2 knockdown is greater in constructs with polyA tracks inserted after the signal peptide, this difference was not significant (P-value <0.05) and was not clearly observed in functional IL2 assay (Fig. 3B-D). However, one can imagine that this discrepancy in the IL2 expression may result from the stalling and frameshifting mechanism of the polyA track.^58^ While the translational regulatory nature polyA track allows for a decrease in gene expression for all constructs, the location may have a differential effect based on factors pertaining to mRNA quality control mechanisms.^59^

By combining our method with clustered regularly interspaced short palindromic repeat (CRISPR)-Cas9-based methods, we can create hypomorphic mutants endogenously. We tested this method by creating endogenous AUF1 and TP53 hypomorphs in diploid HEK293 cell lines and the near-haploid human HAP1, respectively (Fig 3 and 4). The introduction of 12As in the coding sequence of TP53 led to a decrease of ∼40% in both mRNA stability and protein expression. The endogenous insertion of 36As in the gene locus of the *TP53* led to an approximate 90% reduction in mRNA stability, TP53 protein expression was nearly undetectable by Western blot (Fig 3). After inserting polyA tracks of 12 and 18As in the coding region of the AUF1, we saw a reduction in the level of gene expression of approximately 40 and 60%, respectively. Protein expression of the four major isoforms (p37^AUF1^, p40 ^AUF1^, p42^AUF1^, p45^AUF1^) showed similar levels of decrease in protein in expression (Fig. 4). These results are consistent with previously observeded eukaryotic reporter genes polyA track insertions of 12 and 36As.^19^ The ∼40% knockdown in both TP53 and AUF1 in both haploid and diploid cells, respectively, with the insertion of 12As in each gene supports the predictability and effectiveness of this tool various genes and cell types. By creating TP53 hypomorphs in the HAP1 line, we examined the effects of polyA track insertion on TP53 expression by mutating a single allele. The ability to create monoallelic hypomorphs using CRISPR/Cas9 and polyA tracks allows for a feasible and straightforward method for studying phenotype when the expression of one allele is decreased. ^60^ Moreover, TP53 hypomorphs can be used to create a haploid state in haploid cells, such as HAP1, in which diploidization can occur. ^60,61^The use of our titratable method for hypomorph creation may be able to maintain the haploid state through a P53 reduction without promoting G1 cell cycle arrest and apoptosis that sometimes occurs with the deletion of P53.^62^

In creating titratable CRISPR/Cas9 engineered endogenous hypomorphs, we examined the application of this tool in understanding the role of hypomorphic gene expresson of *AUF1* and *TP53* on the expression of their downstream targets. After observing the differential effects of the addition of 12 or 18As on the downregulation of AUF1, we assayed the effect of AUF1 protein reduction on previously defined targets, including *MAT1A, APP, TOP2A, USP1*. ^25,25,63^ Because AUF1 has been demonstrated to either stabilize or destabilize various targets, we expected that reduction in AUF1 protein would have differential effects on the expression of target mRNAs depending on their previously established relationships.^64^ For example, siRNA-mediated silencing of *AUF1* has shown to stabilize the expression of *MAT1A*.^24^ Our results support this relationship between *AUF1* and *MAT1A*. Reduced expression of AUF1 in both 12A and 18A hypomorphs lead to a moderate increase in *MAT1A* transcript expression slightly above wild-type levels (Fig. 4D). Similarly, PAR-CLIP analysis identified the post-transcriptional gene regulation of *APP, TOP2A*, and *USP1* by AUF1.^25^ AUF1 was found to interact with each mRNA through binding sites in the 3’UTR and, together with HuR protein, regulates translational efficiency and stability of target mRNAs. Our results show that the reduction in mRNA stability of each of these genes is related to the level of AUF1 downregulation, with a stepwise decrease in expression of these targets in 12A and 18A hypomorphs, respectively.

We determined the differential expression of TP53 against several cancer-related genes, including *BRCA1, DUSP6, DDIT4, MGMT* and *P21* (Fig. 6). Most notably, we observed that patterns of decreased TP53 expression do not necessarily correlate with the expression levels for downstream targets regulated by TP53. The differential effects of TP53 hypomorohs on DDIT4 and DUSP6 proteins is more than intriguing given that both of these proteins are connected with TP53 status, mTOR pathway, cell metabolism and proliferation.^45^ Also there were relatively stable levels of BRCA1 expression in both cells containing 12A or 36A insertions, while MGMT and P21 were decreased by more than 85% in both cell lines. A larger downregulation of MGMT and P21 than the level of TP53 expression has been supported by previous studies.^49,60,65^ Our study allows for a more quantitative and tunable approach to investigating TP53 expression levels’ role in the modulation of downstream targets previously accomplished with popular methods such as RNAi.^60^ The P21 levels under hydroxyurea treatment in TP53 polyA track engineered cells showed delay in induction compared to wild-type TP53 cells (Fig. 6C).^50^ In the case of MGMT, our tool’s precision and predictability may have utility for uncovering a more cohesive role of TP53 downregulation on MGMT expression and how this regulation affects chemoresistance in certain cancers (Fig. 6D) ^49^.

The utility of hypomorphic mutants created through polyA insertion can be seen in studies elucidating the function of proteins expressed in pathogens such as *Candida albicans*. A recent study in the fungal pathogen *C. albicans* showed a decrease in protein abundance with the insertion of polyA tracks of different lengths in the lanosterol demethylase (Erg11p) and Penta-functional AROM polypeptide (Aro1p), both common targets of antifungals. ^20^ The reduction of both genes’ protein expression using this method was predictable, with the insertion of six or more AAA codons causing a reduction in gene expression. The use of hypomorphic mutants created through polyA tracks in the prevalent pathogen, *C. albicans*, supports this tool’s use in pathogenic microbes. With this tool, microbiologists can investigate the phenotypic effects of genetic manipulation in host organisms without the use of xenobiotics and external environment changes. The results of this study and previous studies showing the effects of polyA track gene insertion undoubtedly support this tool’s utility. The tool has been shown to be effective in both eukaryotic and prokaryotic unicellular and multicellular organisms utilizing both reporter constructs and CRISPR/Cas9. We predict the use of CRISPR/Cas9 for introduction of polyA tracks and generation of titratable and straightforward hypomorphic mutants that will prove invaluable for synthetic biology and genetic studies.

## MATERIALS AND METHODS

### Cell Culture

Hap1 TP53 cells were cultured in IMDM media and supplemented with 10% Fetal-GRO and 5% penicillin and streptomycin (Gibco) and L-glutamine (Gibco). Flp-In T-REx 293 cells AUF1 cell line was cultured in Dulbecco’s modified Eagle’s medium (DMEM) (Gibco) and supplemented with 10% Heat-Inactivated Fetal Bovine Serum (Gibco), 5% minimum essential medium nonessential amino acids (100 ×, Gibco), 5% penicillin and streptomycin (Gibco) and L-glutamine (Gibco).

Additionally, 5 μg ml−1 of blasticidin and 100 μg ml−1 of Zeocin was added to media for cell line maintenance. Flp-In T-REx 293 cells were cultured in Dulbecco’s modified Eagle’s medium (DMEM) (Gibco) and supplemented with 10% Fetalgro® Bovine Growth Serum (BGS), 5% minimum essential medium nonessential amino acids (100 ×, Gibco), 5% penicillin, streptomycin (Gibco), and L-glutamine (Gibco). Additionally, 5 μg ml−1 of blasticidin and 100 μg ml−1 of Zeocin was added to the media for cell line maintenance. CHO-K cells were cultured in F-12K Medium (Kaighn’s Modification of Ham’s F-12 Medium) (Gibco) supplemented with 10% Fetalgro® Bovine Growth Serum (BGS), 5% minimum essential medium nonessential amino acids (100 ×, Gibco), 5% penicillin and streptomycin (Gibco), and L-glutamine (Gibco).

### DNA Constructs

IL2 and CD20 constructs were created through PCR amplification from cDNA and genomic DNA respectively.PCR products were purified using the Zymoclean Gel DNA Recovery Kit (Zymo Research) and cloned into the pEntryD-Topo vector. From the pEntryD-Topo vector, constructs were subcloned into pcDNA-DEST40 or pcDNA-DEST53 vector for expression using LR clonase recombination (Thermo Fisher Scientific). Constructs were transformed using One Shot TOP10 Chemically Competent E. coli cells (Thermo Fisher Scientific). Sequences for plasmid DNA were confirmed by Sanger sequencing (Genewiz).

### CRISPR/Cas9-mediated Gene Editing

CRISPR/Cas9 genomic editing was performed by the Genome Engineering and IPSC Center (GEIC) at Washington University. sgRNAs were designed according to sequences for both TP53 and AUF1, respectively, acquired from the Ensembl genetic database. sgRNAs were selected based on off-target analysis performed by GEIC-specific algorithms. RNPs containing recombinant CRISPR/Cas9 and sgRNA were transfected to human cell lines using a Nucleofector (Lonza). Stable cell lines were generated by selection of hygrogmycin resistance. After assessment of Cas9 NHEJ nuclease activity by the Cel-1 assay cell lines were proprogated and assessed for positive insertion of polyA tracks by qRT-PCR and sequencing analysis (Supplementary Fig. 1 and 2). A more detailed description of these methods can be found in the recent publication from GEIC.^66^

### Live Imaging and Flow Cytometry of CD20 expressing cells

For live imaging CHO-K cells were electroporated with a mCherry reporter or either wild-type CD20 or CD20 constructs containing an insertion of 12 and 18As, respectively, by the Neon Transfection System(Thermo Fisher Scientific) using the 100 ul tips and cell line-specific protocol (https://www.thermofisher.com/us/en/home/life-science/cell-culture/transfection/neon-transfection-system/neon-transfection-system-cell-line-data.html). 24 hours post-electroporation, cells were washed twice with 1x PBS and blocked for 30 minutes with PBS/BSA buffer (phosphate buffered saline pH 7.4 and 1% BSA). After blocking, cells were gently washed with 1x PBS before being incubated for one hour in 1:500 diulution of FITC-conjugated CD20 antibody (Anti-Human CD20 FITC 2H7 – Thermo) in PBS/BSA buffer. Following incubation, cells were washed with 1x PBS prior to and after incubation with for 1 hour in with a 1:8000 dilution of Hoescht 33342 (Invitrogen)) and five minutes with a 1:5000 dilution of CellMask™ Orange Plasma membrane Stain (Invitrogen) in 1x PBS Cells were washed twice with 1x PBS prior to imaging. Live flourescence imaging was performed at the Washington University Center for Cellular Imaging (WUCCI). Cells were imaged using a using the Nikon A1Rsi scanning confocal microscope in widefield mode using a Nikon CFI Plan Apochromat 20× / 0.75 objective. Emission was filtered at 450±50 nm (DAPI), 525±25 nm (eGFP), and 595±25 nm (mCherry/mTangerine). Cells were maintained at 37 °C with 5% CO2, controlled by a stagetop microscope incubation system (INU TIZW, TOKAI HIT, Japan).

For flow cytometry experiments CHO-k cells were seeded at a density of 1.0*10^6 cells in a six-well plate 24 hours before transfection. Cells were cotransfected with mCherry reporters and either wild-type CD20 or CD20 constructs containing insertion of 12 or 18As, respectively. 48 hours post transfection cells were trypsinized and pelleted. Cell pellets were washed twice with 1x PBS before being blocked with PBS/BSA buffer for 30 minutes on ice. After blocking, cell pellets were washed once with 1x PBS before a 1hour incubation with a 1:500 dilution of FITC-conjugated CD20 antibody. Cell suspensions were filtered with (Partec North America CELLTRICS 100UM NS FLTR). Cytofluorimetric analysis was performed to enumerate CD20 positive and mCherry positive CHO-k cells using the BD FACSCalibur sytem (BD Biosciences).

### Enzyme-Linked Immunosorbent Assay

Flp-In T-REx 293 cells were plated at a density of 1.5×10^6 cells in one well of the six-well plates. Twenty-four hours after transfection with WT IL2 or IL2 constructs containing inserted polyA tracks media was collected. Media from was centrifuged for 1,500 rpm centrifuge at 4°C for 10 mins, and cell supernatant was collected. The supernatant was used immediately or stored at −80°C. The solid-phase sandwich ELISA (enzyme-linked immunosorbent assay) (Thermo Fisher, EH2IL2) was used to measure IL2 concentration from cell supernatant according to the manufacturer’s protocol(https://assets.thermofisher.com/TFS-Assets/LSG/manuals/MAN0014639_EH2IL2_ELISA_Kit_PI.pdf). Cell supernatant was diluted with complete cell media at a ratio of 1:10 to be within the concentration of a range of the standard curve. Diluted cell supernatant was further diluted to obtain 1:2,1:4,1:8, and 1:16 dilutions using complete media. Absorbance measurements were taken using the Synergy H4 Hybrid Multi-Mode Microplate Reader (Bioteck) at 450 minus 550 nm. Concentrations were interpolated from standard curves created with GraphPad Prism statistical software.

### IL2 Luciferase Bioassay

According to the manufacturer’s protocol, cell supernatant from untransfected and cells transfected with IL2 constructs was diluted 1:12 with complete media and used to perform the IL2 Bioassay (Promega, JA2201). Luminescence (RLU) was measured using the GloMax® Explorer Multimode Microplate Reader. Average RLU values were substracted from background wells. Relative Luciferase expression was measured in reference to samples from cells transfected with wild-type IL2 constructs.

### Hydroxyurea treatment

Hydroxyurea (Sigma Aldrich) was kindly provided by Dr. Zhongsheng You. WT HAP1, HAP1 TP53 12A and HAP1 TP53 18A cells, were transferred into 12 well plate 24 hours before the hydroxyurea treatment. Hydroxyurea was added in the media to final concentration of 1.5 mmol. Non-treated and hydoxyurea-treated cells were collected 3 and 24 hours after addition of hydroxyurea. Cell lysates were used for western blot analysis.

### Western blot analysis

Total cell lysates for Western blot analysis of TP53, AUF1, and CD20 hypomorphs were prepared with passive lysis buffer (Promega) and homogenized by passing five times through a 25-gauge needle and syringe. Total protein was measured from cell lysis using the Bio-Rad DC Protein Assay according to the manufacturer’s instructions(http://www.bio-rad.com/webroot/web/pdf/lsr/literature/LIT448.pdf). CD20 Samples were prepared in a 1x Bio-Rad XT sample buffer with 5% β-mercaptoethanol with equal total protein concentrations. Samples were boiled at 55 degrees Celcius for 10 minutes before loading on SDS-page gel. AUF1 and TP53 samples were prepared in a 1x Bio-Rad XT sample buffer with a 1x XT reducing agent (Biorad). Samples were boiled at 95 degrees Celcius for 5 minutes before loading on SDS-page gel.

The CD20 blot was blocked overnight in 5% milk (w/v) in 1 × phosphate-buffered saline with 0.1% Tween 20 (PBST). Blots were incubated in a 1:500 dilution of CD20 primary antibody overnight (Thermo Fisher Scientific PA5-16701) and a 1:5000 dilution of anti-GFP (Takara, 632381). Blots were washed (3 × 10 mins) in TBST before and after incubating with a 1:10,000 anti-mouse secondary antibody (Cell Signaling, 7076S) overnight. The blots were washed overnight before being incubated with a 1:5000 dilution of anti-actin-HRP for two hours at room temperature (Cell Signaling, 13E5).

The TP53, BRCA1, DUSP6, DDIT4, and MGMT blots were blocked overnight in 5% milk (w/v) in 1 × phosphate-buffered saline with 0.1% Tween 20 (PBST). All blots except for MGMT (incubated for 1.5 hrs at room temperature) were incubated overnight at 4 degrees Celcius in a 1:1000 dilution with the following primary antibodies: anti-TP53 (Cell Signaling, 2524), anti-BRCA1 (Abclonal, A11034), anti-DUSP6 (Santa Cruz, 377070), anti-DDIT4 (Proteintech, 10638-1-AP), P21 (Cell Signaling, 2947)and MGMT (Novus Biologicals, NB-100-168). Blots were washed (3 × 10 mins) in TBST before and after incubating with either a 1:10,000 anti-mouse secondary antibody (Cell Signaling, 7076S) or 1:10,000 anti-rabbit secondary antibody (Cell Signaling, 7074S) overnight. The blots were mildly stripped, washed vigorously, and blocked overnight in 5% milk (w/v) in 1 × phosphate-buffered saline with 0.1% Tween 20 (PBST) before being incubated with a 1:5000 dilution of anti-GAPDH-HRP (Biolegend, 649203) or 1:2000 dilution of anti-ACTIN-HRP (5125)

The AUF1 blots were blocked overnight in 5% milk (w/v) in 1 × phosphate-buffered saline with 0.1% Tween 20 (PBST). The blots were incubated overnight at 4 degrees Celcius in a 1:2000 dilution of anti-AUF1 antibody (Cell Signaling D6O4F). The blots were washed (3 × 10 mins) in TBST before and after incubating with a 1:10,000 anti-rabbit secondary antibody (Cell Signaling, 7074S). The blots were washed overnight before being incubated with a 1:5000 dilution of anti-HSP70-HRP (need catalog number) for one hour at room temperature. All blots were washed 3 × 10 mins in TBST before being developed with SuperSignal™ HRP substrate (Thermofisher Scientific). Images for all blots were generated by the Bio-Rad Molecular Imager ChemiDoc XRS System with Image Lab software.

### RNA isolation and cDNA synthesis

Total RNA was extracted from Flp-In T-REx 293, CHO-K, and HAP1 cell pellets using the illustra triplePrep Kit (GE Healthcare Life Sciences) according to the manufacturer’s instructions. An optional on-column DNAse treatment was performed to remove genomic DNA(https://cdn.gelifesciences.com/dmm3bwsv3/AssetStream.aspx?mediaformatid=10061&destinationid=10016&assetid=13444). RNA concentration was measured by NanoDrop (OD260/280). One microgram of total RNA was used to synthesize cDNA using the SuperScript IV VILO Master Mix (Thermofisher Scientific).

### qPCR analysis

qPCR was performed with iQ SYBR Green Supermix (Bio-Rad) in the Bio-Rad CFX96 Real-Time System using Bio-Rad CFX Manager 3.0 software. The IL2 and CD20 constructs were identified using the following forward primer: 5′-CCCAAGCTGGCTAGTTAAGC-3′ and reverse primers: 5′-GGTTTTGGACCAGATTGCAT-3′ and 5′-TCCAGCAGTAAATGCAGT-3′, respectively. All other genes in this study were identified using gene-specific primers.

### Statistics

Statistical significances between control and experimental groups was calculated using unpaired Student’s t-tests using Graph Pad Prism version 9.1.2 (GraphPad Software, San Diego, California USA). Analysis were performed seperately for each experiment.

## ACKNOWLEDGMENTS

We thank Dr. Caitlin Hanlon and members of Djuranovic’s lab for helpful comments. This work is supported by NIH R01 GM112824, NIH R01 GM136823, NIMH R01 MH116999 and Siteman investment program funds to SD and SPD, and NIH T32 GM: 007067 and R25HG00668708 to GP. We are also thankful to Ratner’s Lab, GeIC and WUCCI centers of Washington University on the help with this project. The Washington University LEAP Gap Fund supported this project through the Skandalaris Center for Interdisciplinary Innovation and Entrepreneurship under award number #1014, the Washington University Institute of Clinical and Translational Sciences, and the NIH/National Center for Advancing Translational Sciences (NCATS) grant UL1TR002345. PolyA tracks technology is part of U.S. and international Patent, Serial No. PCT/US2017/041766.

## AUTHOR CONTRIBUTIONS

GP and SPD performed all experiments. GP, SPD and SD performed data analysis and wrote the manuscript. SPD and SD conceived the project and provided funding. All authors were involved in editing the manuscript.

