## Supplementary Figures for "Gene dosage effects of polyA track engineered hypomorphs"

**Supplementary Figure 1. (Powell et al. 2021)**

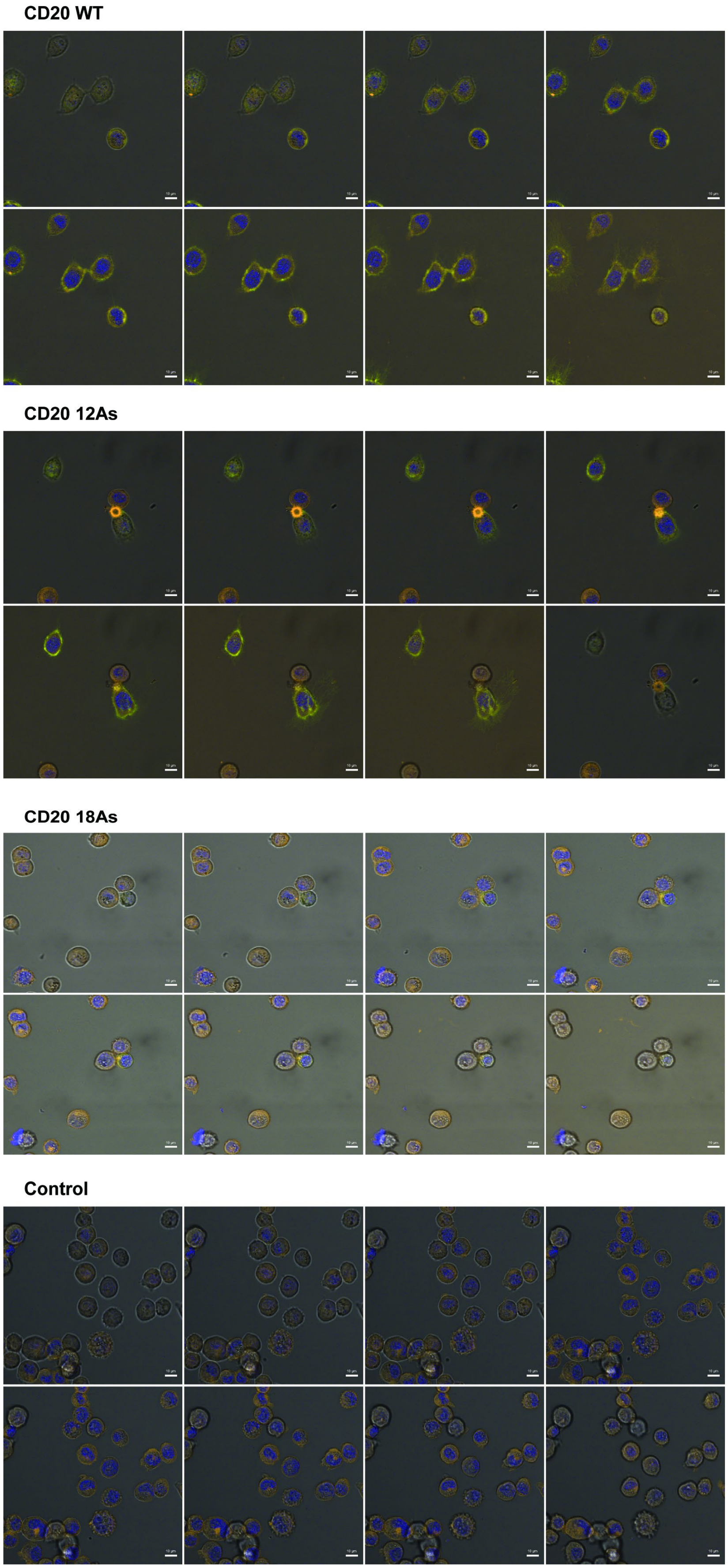


**Supplementary Figure 1. CD20 hypomorphs live imaging.** Consecutive z-stacks of merged fluorescent and bright field microscopy images of live CHO cells expressing CD20, CD20 12As and CD 20 18As constructs. CD20 positive cells were labelled using anti-CD20-2H7 FITC-labelled antibody. The cell membrane is labeled with CellMask™ Orange stain and the DNA was labeled with Hoechst 33342 stain. White scale bar represents 10 um. The control non-transfected CHO cells are indicated.

**Supplementary Figure 2.** (Powell et al. 2021)

**A.**

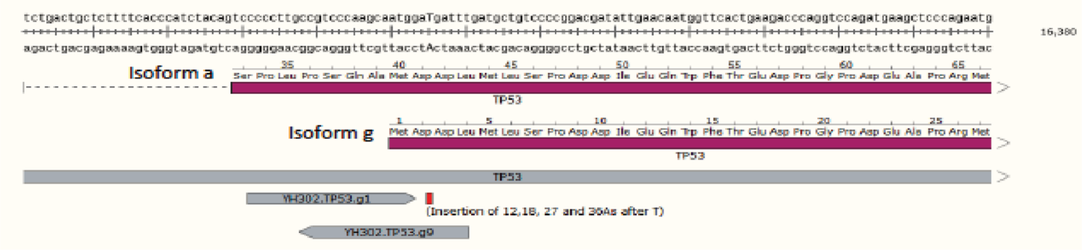


**B.**

| Name | Score | Doench16 | gRNA | long_0 | long_1 | long_2 | long_3 | short_0 | SNP | SNP_count | BP | Distance_ |
|---|---|---|---|---|---|---|---|---|---|---|---|---|
| EA310.HNRNPD.sp1 | 100 | 0.429 | CTAGTTTCGGTTCGCGGCAGNGG | 1 | 1 | 1 | 1 | 6 | NA | 0 | 73 | -52 |
| EA310.HNRNPD.sp2 | 100 | 0.499 | GTTTCGGTTCGCGGCAGCGGNGG | 1 | 1 | 1 | 1 | 10 | NA | 0 | 73 | -49 |
| EA310.HNRNPD.sp3 | 99 | 0.441 | TTTCGGTTCGCGGCAGCGGCNGG | 1 | 1 | 1 | 2 | 28 | NA | 0 | 73 | -48 |
| EA310.HNRNPD.sp13 | 97 | 0.661 | GTCGGAGGAGCAGTTCGGCGNGG | 1 | 1 | 1 | 8 | 2 | NA | 0 | 73 | 12 |
| EA310.HNRNPD.sp6 | 97 | 0.523 | GGGTGTAGTCTCGGCGGCAGNGG | 1 | 1 | 1 | 8 | 6 | NA | 0 | 73 | -28 |
| EA310.HNRNPD.sp11 | 96 | 0.625 | ATGTCGGAGGAGCAGTTCGGNGG | 1 | 1 | 1 | 14 | 13 | NA | 0 | 73 | 10 |
| EA310.HNRNPD.sp5 | 95 | 0.721 | AGCGGCGGGTGTAGTCTCGGNGG | 1 | 1 | 1 | 7 | 6 | NA | 0 | 73 | -34 |
| EA310.HNRNPD.sp16 | 93 | 0.372 | GGAGCAGTTCGGCGGGGACGNGG | 1 | 1 | 1 | 11 | 10 | NA | 0 | 73 | 18 |
| EA310.HNRNPD.sp17 | 85 | 0.431 | GCAGTTCGGCGGGGACGGGGNGG | 1 | 1 | 1 | 32 | 126 | NA | 0 | 73 | 21 |
| EA310.HNRNPD.sa1 | NA | NA | TAGTTTCGGTTCGCGGCAGCGNNGRRT | 1 | 1 | 1 | 1 | 1 | NA | 0 | 73 | -50 |
| EA310.HNRNPD.sp4 | 98 | 0.481 | GGCAGCGGCGGGTGTAGTCTNGG | 1 | 1 | 2 | 9 | 5 | NA | 0 | 73 | -37 |
| EA310.HNRNPD.sp8 | 91 | 0.651 | CGGAGACACTAGCACTATGTNGG | 1 | 1 | 2 | 12 | 31 | NA | 0 | 73 | -5 |
| EA310.HNRNPD.sp7 | 87 | 0.543 | TGTAGTCTCGGCGGCAGCGGNGG | 1 | 1 | 2 | 20 | 400 | NA | 0 | 73 | -25 |
| EA310.HNRNPD.sp30 | 81 | 0.45 | GGCTCGGCGGGCGAGCAGGANGG | 1 | 1 | 2 | 34 | 47 | NA | 0 | 73 | 70 |
| EA310.HNRNPD.sp15 | 95 | 0.335 | AGGAGCAGTTCGGCGGGGACNGG | 1 | 1 | 3 | 13 | 4 | NA | 0 | 73 | 17 |
| EA310.HNRNPD.sp14 | 89 | 0.403 | GAGGAGCAGTTCGGCGGGGANGG | 1 | 1 | 3 | 29 | 7 | NA | 0 | 73 | 16 |
| EA310.HNRNPD.sp28 | 77 | 0.424 | AGGCGGCTCGGCGGGCGAGCNGG | 1 | 1 | 3 | 27 | 28 | NA | 0 | 73 | 66 |
| EA310.HNRNPD.sp27 | 94 | 0.375 | GCGGCGGTAGGCGGCTCGGCNGG | 1 | 1 | 4 | 21 | 12 | NA | 0 | 73 | 58 |
| EA310.HNRNPD.sp9 | 88 | 0.688 | AGACACTAGCACTATGTCGGNGG | 1 | 1 | 4 | 12 | 11 | NA | 0 | 73 | -2 |
| EA310.HNRNPD.sp26 | 87 | 0.468 | GGCGGCGGTAGGCGGCTCGGNGG | 1 | 1 | 4 | 25 | 6 | NA | 0 | 73 | 57 |
| EA310.HNRNPD.sp24 | 76 | 0.441 | GCGGCAACGGCGGCGGTAGGNGG | 1 | 1 | 5 | 60 | 34 | NA | 0 | 73 | 49 |
| EA310.HNRNPD.sp18 | 61 | 0.353 | GTTCGGCGGGGACGGGGCGGNGG | 1 | 1 | 5 | 64 | 100 | NA | 0 | 73 | 24 |
| EA310.HNRNPD.sp20 | 39 | 0.328 | CGGGGCGGCGGCAGCGGCAANGG | 1 | 1 | 5 | 201 | 19 | NA | 0 | 73 | 36 |
| EA310.HNRNPD.sp32 | 41 | 0.572 | CGAGCAGGAGGGAGCCATGGNGG | 1 | 1 | 7 | 110 | 257 | NA | 0 | 73 | 81 |

**C.**

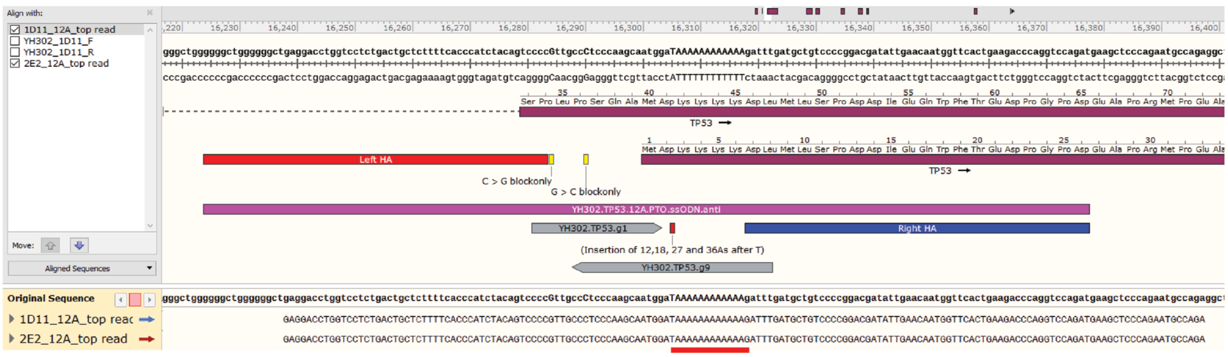


**D.**

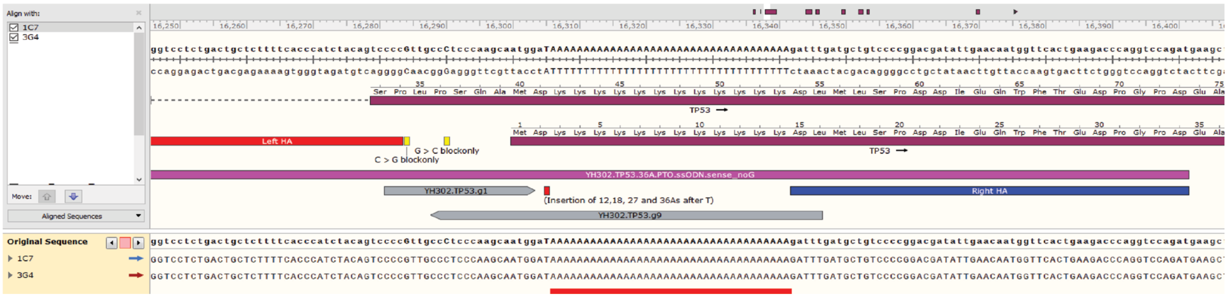


**Supplementary Figure 2. CRISPR/Cas9 design validation for endogenous polyA insertion in the TP53 gene locus of HAP1 cells. A.** Design schema shown with both coding and translated sequence representing the location of specific gRNAs and polyA track insertion. Poly insertion represented by red rectangle. **B.** OFF-target analysis for gRNAs. Selected gRNAs along with number of genomic mismatches to longer and shorter regions of the selected target sequence (long_0,long_1, long_3,short_0) are highlighted in red. SNP (Single nucleotide polymorphism), SNP count, and distance from cut site in base-pairs (BP) are also shown. **C.** Sequencing results for 12A and 36A single clones, respectively. Results are shown for 2 clones for each insert (12 or 36A), both in the forward direction. In addition to target sequence and gDNAs being represented, the location of the ssODN (fuschia) in shown with both 5' and 3' homology arms (red and blue, respectively). The sequenced results for the region surrounding the polyA track insertion (red underline) is shown.

**Supplementary Figure 3.** (Powell et al. 2021)

**A.**

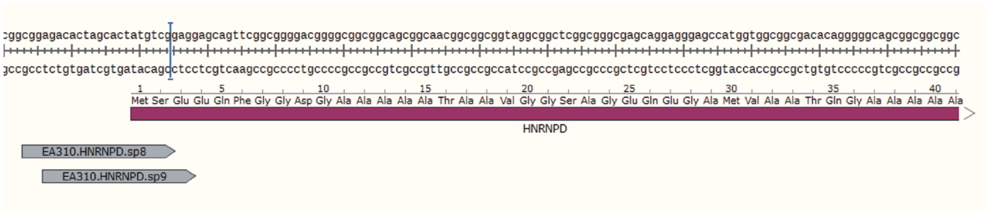


**B.**

| Name | Score | Doench16 | gRNA | long_0 | long_1 | long_2 | long_3 | short_0 | SNP | SNP_count | BP | Distance_fi |
|---|---|---|---|---|---|---|---|---|---|---|---|---|
| EA310.HNRNPD.sp1 | 100 | 0.429 | CTAGTTTCGGTTCGCGGCAGNGG | 1 | 1 | 1 | 1 | 6 | NA | 0 | 73 | -52 |
| EA310.HNRNPD.sp2 | 100 | 0.499 | GTTTCGGTTCGCGGCAGCGGNGG | 1 | 1 | 1 | 1 | 10 | NA | 0 | 73 | -49 |
| EA310.HNRNPD.sp3 | 99 | 0.441 | TTTCGGTTCGCGGCAGCGGCNGG | 1 | 1 | 1 | 2 | 28 | NA | 0 | 73 | -48 |
| EA310.HNRNPD.sp13 | 97 | 0.661 | GTCGGAGGAGCAGTTCGGCGNGG | 1 | 1 | 1 | 8 | 2 | NA | 0 | 73 | 12 |
| EA310.HNRNPD.sp6 | 97 | 0.523 | GGGTGTAGTCTCGGCGGCAGNGG | 1 | 1 | 1 | 8 | 6 | NA | 0 | 73 | -28 |
| EA310.HNRNPD.sp11 | 96 | 0.625 | ATGTCGGAGGAGCAGTTCGGNGG | 1 | 1 | 1 | 14 | 13 | NA | 0 | 73 | 10 |
| EA310.HNRNPD.sp5 | 95 | 0.721 | AGCGGCGGGTGTAGTCTCGGNGG | 1 | 1 | 1 | 7 | 6 | NA | 0 | 73 | -34 |
| EA310.HNRNPD.sp16 | 93 | 0.372 | GGAGCAGTTCGGCGGGGACGNGG | 1 | 1 | 1 | 11 | 10 | NA | 0 | 73 | 18 |
| EA310.HNRNPD.sp17 | 85 | 0.431 | GCAGTTCGGCGGGGACGGGGNGG | 1 | 1 | 1 | 32 | 126 | NA | 0 | 73 | 21 |
| EA310.HNRNPD.sa1 | NA | NA | TAGTTTCGGTTCGCGGCAGCGNNGRRT | 1 | 1 | 1 | 1 | 1 | NA | 0 | 73 | -50 |
| EA310.HNRNPD.sp4 | 98 | 0.481 | GGCAGCGGCGGGTGTAGTCTNGG | 1 | 1 | 2 | 9 | 5 | NA | 0 | 73 | -37 |
| EA310.HNRNPD.sp8 | 91 | 0.651 | CGGAGACACTAGCACTATGTNGG | 1 | 1 | 2 | 12 | 31 | NA | 0 | 73 | -5 |
| EA310.HNRNPD.sp7 | 87 | 0.543 | TGTAGTCTCGGCGGCAGCGGNGG | 1 | 1 | 2 | 20 | 400 | NA | 0 | 73 | -25 |
| EA310.HNRNPD.sp30 | 81 | 0.45 | GGCTCGGCGGGCGAGCAGGANGG | 1 | 1 | 2 | 34 | 47 | NA | 0 | 73 | 70 |
| EA310.HNRNPD.sp15 | 95 | 0.335 | AGGAGCAGTTCGGCGGGGACNGG | 1 | 1 | 3 | 13 | 4 | NA | 0 | 73 | 17 |
| EA310.HNRNPD.sp14 | 89 | 0.403 | GAGGAGCAGTTCGGCGGGGANGG | 1 | 1 | 3 | 29 | 7 | NA | 0 | 73 | 16 |
| EA310.HNRNPD.sp28 | 77 | 0.424 | AGGCGGCTCGGCGGGCGAGCNGG | 1 | 1 | 3 | 27 | 28 | NA | 0 | 73 | 66 |
| EA310.HNRNPD.sp27 | 94 | 0.375 | GCGGCGGTAGGCGGCTCGGCNGG | 1 | 1 | 4 | 21 | 12 | NA | 0 | 73 | 58 |
| EA310.HNRNPD.sp9 | 88 | 0.688 | AGACACTAGCACTATGTCGGNGG | 1 | 1 | 4 | 12 | 11 | NA | 0 | 73 | -2 |
| EA310.HNRNPD.sp26 | 87 | 0.468 | GGCGGCGGTAGGCGGCTCGGNGG | 1 | 1 | 4 | 25 | 6 | NA | 0 | 73 | 57 |
| EA310.HNRNPD.sp24 | 76 | 0.441 | GCGGCAACGGCGGCGGTAGGNGG | 1 | 1 | 5 | 60 | 34 | NA | 0 | 73 | 49 |
| EA310.HNRNPD.sp18 | 61 | 0.353 | GTTCGGCGGGGACGGGGCGGNGG | 1 | 1 | 5 | 64 | 100 | NA | 0 | 73 | 24 |
| EA310.HNRNPD.sp20 | 39 | 0.328 | CGGGGCGGCGGCAGCGGCAANGG | 1 | 1 | 5 | 201 | 19 | NA | 0 | 73 | 36 |
| EA310.HNRNPD.sp32 | 41 | 0.572 | CGAGCAGGAGGGAGCCATGGNGG | 1 | 1 | 7 | 110 | 257 | NA | 0 | 73 | 81 |

**C.**

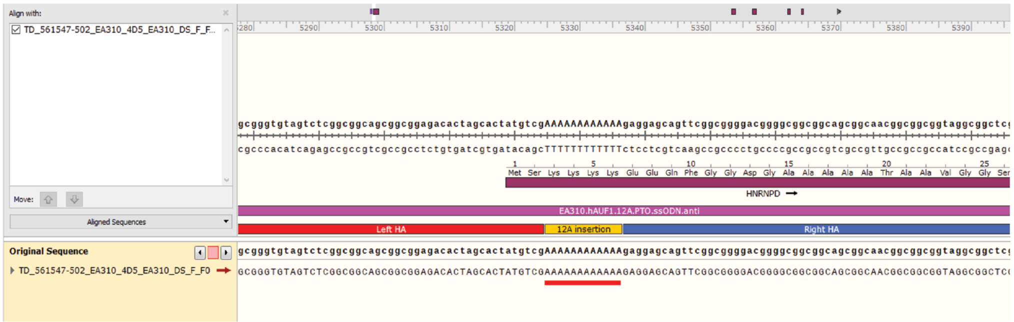


**D.**

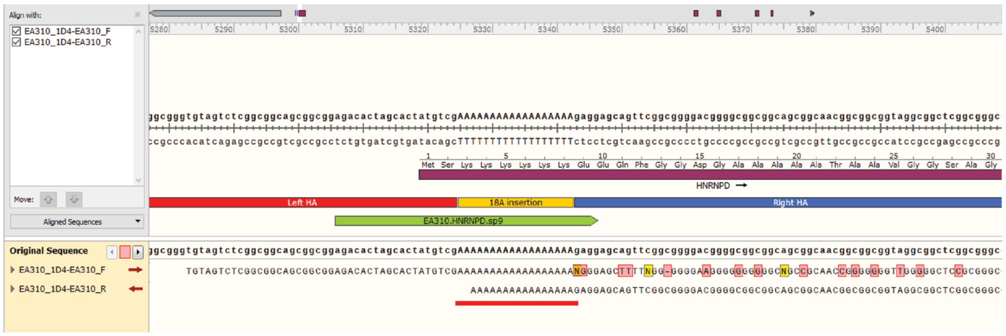


**Supplementary Figure 3. CRISPR/Cas9 Design Validation for Endogenous PolyA insertion in the AUF1 gene locus of HEK293 cells. A.** Design schema shown with both coding and translated sequence representing the location of specific gRNAs and polyA track insertion. Poly insertion represented by red rectangle. **B.** OFF-target analysis for gRNAs. Selected gRNAs along with number of genomic mismatches to longer and shorter regions of the selected target sequence (long_0,long_1, long_3,short_0) are highlighted in red. SNP (Single nucleotide polymorphism), SNP count, and distance from cut site in base-pairs (BP) are also shown. **C.** Sequencing results for a single 12A clone is shown.The target sequence is shown in addition to the location of the ssODN (fuschia) in shown with both 5' and 3' homology arms (red and blue, respectively). The sequenced results for the region surrounding the polyA track insertion (red underline) is shown. **D.** Sequencing results for a single 12A clone in both the forward and reverse directions in shown. The sequenced results for the polyA track insertion (red underline) is shown.
